# Effects of local and landscape factors on paddy field arthropod diversity: from species to communities

**DOI:** 10.1101/2024.01.17.575529

**Authors:** Jia-Ang Ou, Chi-Lun Huang, Chi-Wei Tsai, Chuan-Kai Ho

**Affiliations:** Institute of Ecology and Evolutionary Biology, National Taiwan University, Taipei, Taiwan; Department of Entomology, National Taiwan University, Taipei 106, Taiwan; Department of Life Science, National Taiwan University, Taipei, Taiwan

**Keywords:** Agroecology, Biodiversity, Spatial scales, Trophic guilds, JSDM

## Abstract

Agricultural landscapes are dynamic mosaics where environmental conditions vary across spatial scales. Identifying the spatial scales of environmental variables relevant to biodiversity is therefore essential for safeguarding biodiversity and the ecosystem services they provide. Although species richness is a useful aggregate metric the captures the responses of biodiversity as a whole, few studies have examined how species-specific responses to environmental variables give rise to community-level patterns. Failure to account for the heterogeneity in species responses may undermine potential processes that structure biodiversity. In this study, we sampled paddy-field inhabiting arthropods across the first growing season and examined how local (organic farming, water depth, and crop height) and landscape factors (forest cover and connectivity) affect their abundance and distribution. We built a spatio-temporally explicit joint species distribution model (JSDM) to answer three main questions. 1) How do species differ in their response to local and landscapes factors. 2) Can trophic guild explain the variation in species responses to environmental covariates, and 3) How do species-specific responses scale-up to structure species richness at the community-level? Our results show that the mean importance of environmental covariates was low overall (mean R^2^ = 0.21), suggesting the prevalence of stochasticity in structuring species abundances. However, environmental importance varied substantially across species (highest R^2^ = 0.87 and lowest R^2^ = 0.01) and that trophic guild explained only approximately 15% of species responses, which indicates high idiosyncrasy in species responses both within and among guilds. At the community-level, we found that species richness responds negligibly to local factors while increasing with forest cover at the landscape scale. The importance of each environmental covariate was reflected by the variability of species responses. Lastly, we found no evidence of dispersal limitation in structuring species abundances. Taken together, our study suggests that many paddy field arthropods exhibit transient dynamics and that environmental effects are highly species-specific, even within the same trophic guild. Most importantly, our study highlights the need to understand biodiversity from the species level to understand the processes that structure them in dynamic mosaic landscapes.

## Introduction

### Agriculture and biodiversity: threats and opportunities

To make space for contemporary agriculture, native habitat conversion has become a major driver of biodiversity loss worldwide (Foley et al., 2005; IPBES, 2019). In addition to habitat loss, local agricultural practices such as pesticide and synthetic fertilizer use also pose a threat to the abundance and richness of species occupying these landscapes (Tilman et al., 2001). To meet societal needs and conservation objectives, land managers and researchers have been exploring ways in which human-dominated landscapes can be best managed to protect biodiversity while maximizing production (Kremen & Merenlender, 2018). A potential solution is to utilize the ecosystem services provided by the abundance and richness of biodiversity such as pest control and pollination (Dainese et al., 2019). However, the heterogeneity of environmental variables that vary across spatial scales in agroecosystems make predicting the drivers of biodiversity change challenging (Tscharntke et al 2005). Thus, understanding the spatial scales relevant to farmland biodiversity has been an area of active research in agroecology.

### Local and landscape factors in agroecosystems

Agricultural landscapes are often described as dynamic mosaics which contain a matrix of semi-natural vegetation, ephemeral farmland, freshwater resources, and human settlement (Dominik et al., 2017; Tscharntke et al., 2005; Vandermeer et al., 2010). Because mobile organisms disperse, forage, and sometimes complete their life-cycle stages (Gurr et al., 2017) among these habitats, it is absolutely essential to incorporate both local and landscapes factors to understand the determinants of biodiversity even at the local farmland scale — the scale in which farmers benefit from ecosystem services (Gonthier et al., 2014; Power, 2010; Tscharntke et al., 2005). At local scales for example, farms can differ in their use of synthetic pesticides, fertilizers, variety of crop grown, tillage, and water use. At the landscape scale, farms can differ in their surrounding landscape composition, amount of natural habitat, and connectivity to those habitats and other farms. While many studies have shown that these factors are important for biodiversity, the results are mixed and highly dependent on species and study system. For example, although organic farming can increase the abundance and richness of local farmland biodiversity (Bengtsson et al., 2005; Hole et al., 2005; Letourneau & Bothwell, 2008), these benefits tend to depend on taxonomic group (Fuller et al., 2005), landscape context (Rundlöf & Smith, 2006; Winqvist et al., 2011), and spatial scale (Gabriel et al., 2010; Schneider et al., 2014). These contingent effects of organic farming suggest that species-specific responses and spatial processes are prevalent and may jointly structure local farmland biodiversity.

### Heterogeneity in species responses

Although species richness is a metric commonly used to quantify biodiversity (Magurran, 2004), doing so undermines the potential variability in species responses to environmental factors. In agroecosystems particularly, understanding species specific responses is important because species can vary in their amount and mode of contribution to ecosystem services in the field. For example, parasitoids may provide stronger suppression of crop pests due to their higher specificity relative to generalist predators (Hassell & May, 1986; Schmidt et al., 2003; Snyder & Ives, 2001; but see Symondson et al., 2002 for counter examples). Predator-prey interactions may also exhibit oscillatory dynamics and thus, delivery of pest control services may depend on temporal scale (Chaplin-Kramer et al., 2013). Furthermore, species are likely to vary in their responses to environmental factors due to differences in ecological requirements and dispersal abilities. For instance, broad organismal groups such as plants, invertebrates and vertebrates can differ substantially in their sessility which in turn influence their spatial sensitivity to environmental factors (Gonthier et al., 2014). But even within the same functional group, species can operate at very different scales. For example, within natural enemy species, ballooning spiders may show responses to the percentage of non-crop vegetation at spatial scales beyond 3km (Schmidt et al., 2005) while parasitoids respond to variables at finer scales between 0.5-2km (Thies et al., 2005).

In farmlands, ecological communities are likely to contain co-occurring species that differ substantially in their environmental requirements, dispersal ability, and responses to environment fluctuations. The variability in species responses can explain, at least in part, the inconsistencies in, and system-specific effects of, local and landscape factors on farmland biodiversity (Bengtsson et al., 2005; Fuller et al., 2005; Gabriel et al., 2010; Hole et al., 2005; Karp, Chaplin-Kramer, et al., 2018). Despite the mounting evidence of species-specific responses and the inconsistent effects of local and landscape factors, very few studies have linked these two interrelated components explicitly. Furthermore, although broad functional group classifications help condense information of hyperdiverse communities such as arthropods (Moran et al., 1982), the assumption that species within broad functional groups are similar in their response to environmental variables remains to be tested. As biodiversity necessarily arises from the collective responses of individual species, confronting the heterogeneity in species responses in relation to environmental factors at multiple spatial scales, may facilitate better understanding of how biodiversity is structured in these mosaic landscapes (Frishkoff & Karp, 2019; Tscharntke et al., 2012).

### Study objective

In this study, we take a multi-scale and species-centric approach to disentangle the relative importance of local and landscape factors on farmland biodiversity of irrigated paddy fields in subtropical Taiwan (Fig. 2). Specifically, we sampled arthropods— a taxonomically diverse and functionally rich phylum— across the first growing season, measured local and landscape variables, and built a spatio-temporally explicit, hierarchical Joint Species Distribution Model (JSDM) to address three main questions. 1) How do species differ in their response to local and landscapes factors, 2) Can trophic guilds be used as a good proxy of species responses to environmental variables, and 3) How do species-specific responses scale-up to structure species richness at the community-level?

## Materials and Methods

### Study site description

Our study site is situated in Yuanli township of Miaoli county (24° 33′ 48″ N, 120° 49′ 33″ E), Taiwan—a typical human-dominated landscape encompassing a mosaic of semi-natural vegetation, human settlement, small foothills, ocean coastline, and an abundance of rivers (Fig. 1). We specifically selected 14 rice farms — 7 organic and 7 conventionally-farmed— that reflected the landscape heterogeneity of the region whilst ensuring that conventional and organic farms were not spatially aggregated such to be confounded by our variables of interest. Farms were generally small (0.1-0.37 ha) but typical of farms across Taiwan. Like most lowland areas of the island, the climate is warm, humid, and subtropical, with heavy monsoon rainfalls in the summer months. On average, farmers produce two rice crops a year, with the first cropping season occurring between the months of March and July and the second between August and November. Although farms in the region are still predominately managed under conventional methods, organic farming has been steadily increasing in response to rising environmental awareness, with government subsidies providing incentives to help farmers make the transition. In contrast to conventional farms, organic farms are prohibited from using chemical or synthetic inputs of any kind, including insecticides, herbicides, and fertilizers.

**Figure 1.**
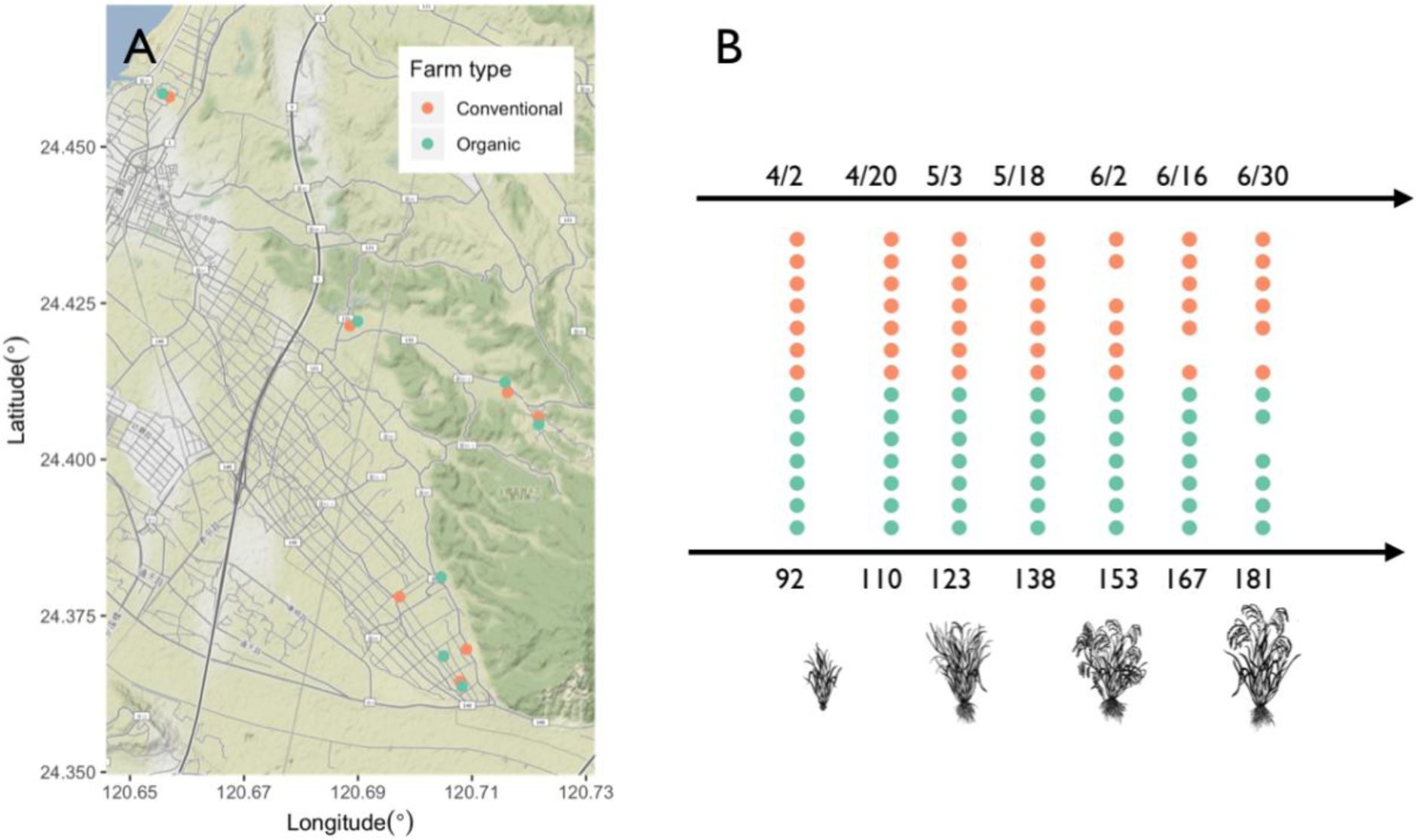
Study site map and sampling scheme. (A) The map shows the 14 farms (points) included in our study system, where colors indicate whether they are conventional (orange) or organic (green). (B) The timeline shows the dates in which the 14 farms (rows) were sampled with Gregorian dates labeled on the top and Julian dates at the bottom. Not all farms could be sampled at certain dates due to access restrictions imposed by the farmers.

**Figure 2.**
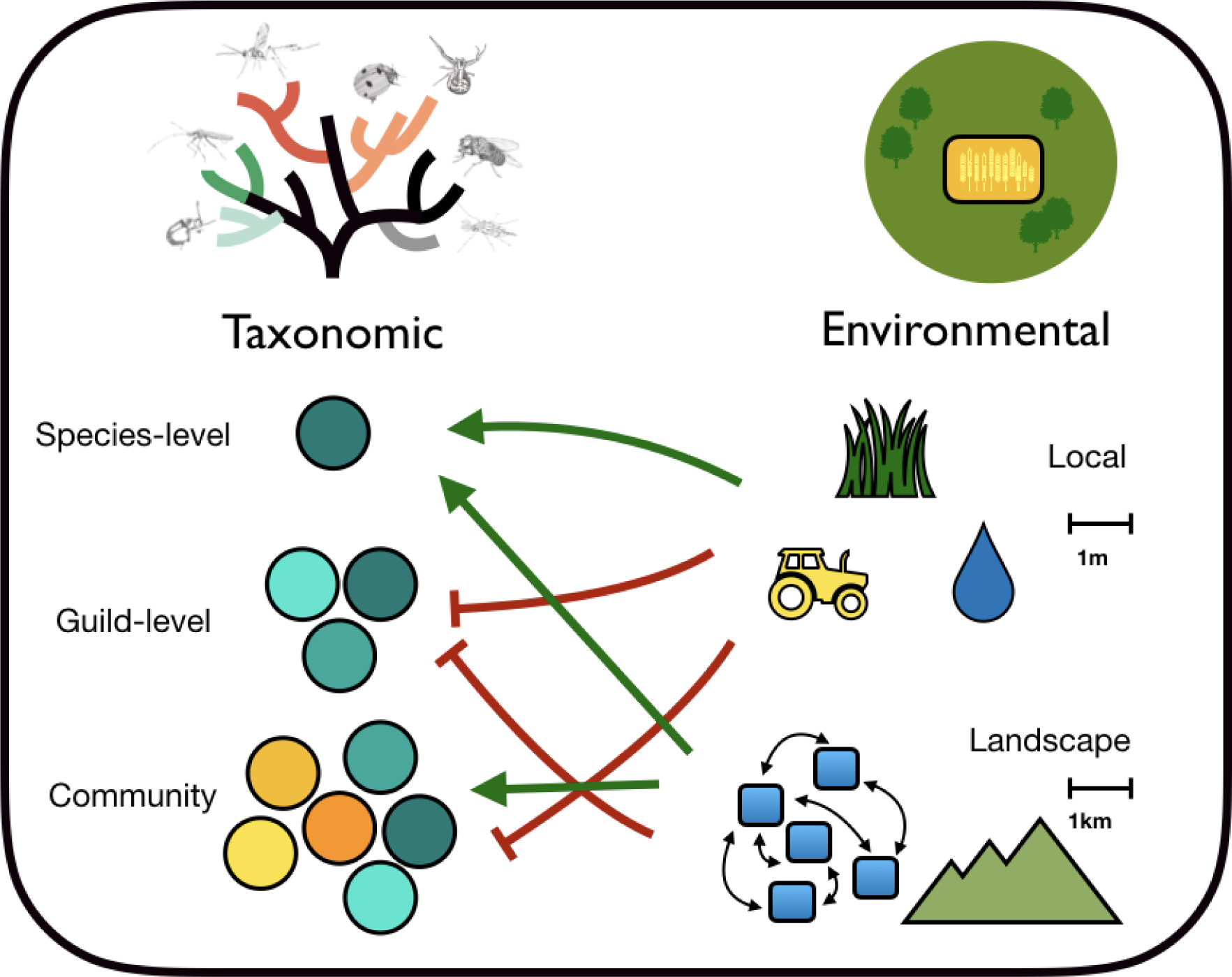
Schematic overview of study objectives. Heterogeneity can arise from the ecological differences (color) among species (circle) and how they respond (color and shape of arrows) to environmental variables that occur at different spatial scales. These different responses to environmental variables may differ depending on the taxonomic scale of observation. For example, local variables may positively affect most species and yet negatively affect community richness if, for instance, the few rare species that contribute to richness differences across sites are negatively affected.

### Arthropod sampling and identification

Arthropod communities were sampled approximately once every two weeks at each farm over the course of the first cropping season (from seedling to ripening; Fig. 1). For standardization purposes and to ensure there was an abundance of arthropods in our samples, sampling events were conducted during the day between 9am to 3pm and only took place when there was at least one day without rain prior to time of sampling. At farmers’ discretion, farms were sometimes inaccessible which led us to lose some data points, resulting in a total of sample size of N = 94. Sweep net samples were conducted by walking along a random transect into the paddy field while sweeping consecutively for 10 times across the crop canopy. Sampled individuals were then immediately placed into ziploc bags and stored without chemical preservatives (e.g., ethanol) at −20°C in the freezer. Following collection, arthropods were counted and. Due to the uncertainty associated with identification at finer taxonomic resolutions, we focus our analysis at the family-level and use the term “family” and “species” interchangeably. We describe our results using the term “family” and reserve “species” for broader use in the discussion section. Although this approach is not ideal, family-level information has been shown to be sufficient in detecting changes in invertebrate community in response to environmental change (Driessen & Kirkpatrick, 2019). Lastly, we assigned trophic guilds to each of the identified family based on published literature (Moran et al., 1982) and our own expert knowledge of the system (Table S1).

### Environmental covariates

Environmental covariates were measured at the local (i.e., farmland) and landscape-level. Local variables include farm type (organic or conventional), crop height, and water depth. Landscape variables include the percentage of surrounding forest cover (or semi-natural vegetation) within 1km radius and the geographic location (given by the geographic coordinates) of each farm. Because crop height and water depth vary with time, these variables were collected in conjunction with arthropod sampling. Specifically, for each sample, crop height and water depth were sampled by haphazardly selecting three rice stems and three locations within paddy fields respectively and taking the average of the three measurements. Crop height was measured as the distance (cm) from the soil to the tip of the rice stem. Water depth was measured as the distance (mm) from the soil to the water surface. Because crop height is strongly correlated with time, we removed the correlation by fitting a simple linear regression across all samples and took the residuals as an index of crop height for all subsequent analyses. Lastly, forest cover was measured by first hand-classifying trees surrounding each farm using satellite images and the polygon tool on Google Earth and processed in Quantum GIS 2.18.7 (QGIS Development team, 2017) to calculate the percentage cover within a 1km radius buffer. Although there are other possible buffer ranges that could be used, we chose 1km because it is standard in many agroecological studies, which makes the result of our study more readily comparable (Gonthier et al., 2014, 2019; Muylaert et al., 2016). All continuous environmental covariates were standardized by subtracting its mean and dividing by its standard deviation for subsequent statistical modeling.

### Data analysis

To examine the effects of environmental covariates on individual species, and in turn, community-level richness, we fitted a Bayesian hierarchical Joint Species Distribution Model (JSDM) using the Hierarchical Modeling of Species Communities (HMSC) framework (Ovaskainen et al., 2017) implemented in the “Hmsc” package ((Tikhonov et al., 2020)) in R. Similar to traditional species distribution models (SDM), JSDM models individual species responses to environmental covariates but has the added advantage over traditional methods by simultaneously accounting for species co-occurrences and hierarchical sampling structures (Ovaskainen et al., 2017; Warton et al., 2015). In this way, the estimated parameters in the model represents the pure environmental effects on species abundances, rather than the combination of environmental effects, biotic interactions, and/or unmeasured environmental covariates. By incorporating spatial information (e.g. spatial coordinates) as random effects, the latent variable approach allows us to estimate the spatial scale (parameter ⍺) in which the residual variation of species abundances operates while capturing the variation among levels of spatial units in the parameter η (Ovaskainen & Abrego, 2020). In other words, the parameter ⍺ describes the extent in which dispersal limitation structures the metacommunity and η describes how species abundances differ among farms. Moreover, simultaneous modeling of multiple species makes the framework more appropriate for modeling sparse ecological data, thereby improving the prediction of rare species by “borrowing” information from the responses of other species (Maguire et al., 2016; Ovaskainen & Soininen, 2011; Zhang et al., 2020). Importantly, the modeling framework can allow predictions to be made at the species, community or trait-level while propagating uncertainty of parameter estimates at the level of the prediction (Ovaskainen et al., 2017).

To examine the effects of environmental covariates on species abundances (i.e. the β parameter), the extent in which these effects can be explained by trophic guild (i.e. a categorical species trait; the γ parameter), and the extent in which dispersal limitation structures the abundance and distribution of arthropods biodiversity, we fitted a spatial-temporally explicit hierarchical JSDM with farm type, crop height, water depth, and forest cover as fixed effects, with sample-level, farm-level (spatially-explicit), and Julian date (temporally-explicit) as nested random effects. Error of the model was assumed to be Poisson distributed. To avoid inflating the importance of environmental covariates, we filtered out extremely rare families which we defined as those that occur in less than 3% of all samples, resulting in a total of 63 families in our analysis.

Parameters of the model were estimated by running 2 MCMC chains, each containing 200,000 iterations, a burn-in length of 1,000, and thinning rate of 100, retaining a total of 4,000 (2 chains * 2000 iterations) posterior samples. Model convergence was assessed by visually inspecting trace plots and the Gelman-Rubin convergence parameter (Gelman & Rubin, 1992). For interpretation of parameter estimates and visualization purposes, we considered estimates in which the 90% posterior credible interval did not overlap zero as having a significant effect. Explanatory power was assessed using Pseudo-R^2^, calculated as the squared Pearson correlation between observed and predicted species abundances. Variance partitioning was used to evaluate the importance of environmental covariates on explaining the variation in species abundances. To visualize the dis/similarity between species in their responses to environmental covariates, we conducted a principal component analysis on the mean of posterior β estimates of each species.

We randomly subsampled 1000 posterior samples for subsequent analysis of posterior communities. To assess the effects of each environmental covariates on the “average species”, we averaged the posterior parameter estimates across species. In a similar fashion, variability in species responses was assessed by taking the standard deviation of responses across species (Frishkoff & Karp, 2019). Effect sizes and significance between covariates were evaluated by taking the mean across the 1000 posterior means/SDs and calculating their 90% highest density interval (HDI). Lastly, to assess the effects of environmental covariates on community richness, emerging from the responses of individual species, we used the posterior parameter distribution to predict species richness (i.e., by taking the row sums of the 1000 predicted posterior species co-occurrences) at each of the original N = 94 sites. The posterior median (rounded to the nearest integer) of species richness at each site was then analyzed by fitting a Poisson-GLMM to examine the effects of environmental covariates on species richness with Julian date as a random effect and the posterior standard deviation as prior weights to account for the uncertainty around predictions (Echeverri et al., 2019; Karp, Frishkoff, et al., 2018; Tingley & Beissinger, 2013). Residual plots were assessed to ensure model assumptions were conformed and test of significance was evaluated using the Wald’s Chi-squared test. We estimated uncertainty of parameter estimates, defined as the 95% confidence interval, using parametric bootstrap (n = 1000). To assess model fit and importance of individual fixed effects, we calculated marginal and semi-partial R^2^ (Jaeger et al., 2017).

## Results

### Arthropod abundance and taxonomic coverage

Across all 94 samples, we identified a total number of 21,833 individuals which belonged to a total of 63 families (Figure S1). Each of the 63 families was assigned into one of 8 distinct trophic guilds: tourist, sap-sucker, predator, parasitoid, leaf-chewer, scavenger, filter-feeder, and unknown. In terms of abundance, the tourist guild was the most abundant, containing a total of 17,993 individuals followed by sap-sucker (2,498) and predator (899). In terms of family richness, the predator guild was richest (19) followed by parasitoid (11) and tourist (10). There was a large variation in the total abundance across families. The Chironomidae family was the most abundant with 13,062 individuals, followed by Ephydridae (4509), and Cicadellidae (1316). The least abundant families were Encrytidae, Linyphiidae, Lycosidae, and Trichogrammatidae, with 5 individuals each. The mean community abundance and mean community richness across all samples were 232.27 and 10.87 respectively.

### Importance of environmental conditions

Assessment of model fits and variance partitioning showed that environmental covariates were of low importance overall (mean R^2^ = 0.20). The most important environmental factor was forest cover (R^2^= 0.031), followed by water depth (R^2^= 0.026), crop height (R^2^= 0.021), and farm type (R^2^= 0.010; Fig. 3). Mean variance of species abundances explained by spatial autocorrelation was 1.72%. Explained variance for sample-level and time was 6.67%, and 3.15% respectively (Fig. 3). At the family-level, explanatory power was highest for Chironomidae (R^2^ = 0.87) and lowest for Staphylinidae (R^2^ = 0.01). For specific covariates, forest cover alone explained up to 15.3% and 13.7% of variation in abundance of Cicadellidae and Coccinellidae respectively (Fig. 3). Water depth explained 15.0% and 14.8% for Thomisidae and Pentatomidae respectively. Crop height explained 8.8% and 7.3% for Thripidae and Araneidae respectively. Lastly, farm type explained 4.9%, 3.5% for Pipunculidae and Acrididae respectively. Taken together, these result show that surrounding landscape is the most important variable in determining the abundance of arthropods inhabiting paddy farms. Although explanatory power and strength of individual fixed effects were low overall, there was large discrepancy among species in the extent to which environmental covariates were able to explain the variation in species abundances, indicating that the importance of environmental condition varied across taxa.

**Figure 3.**
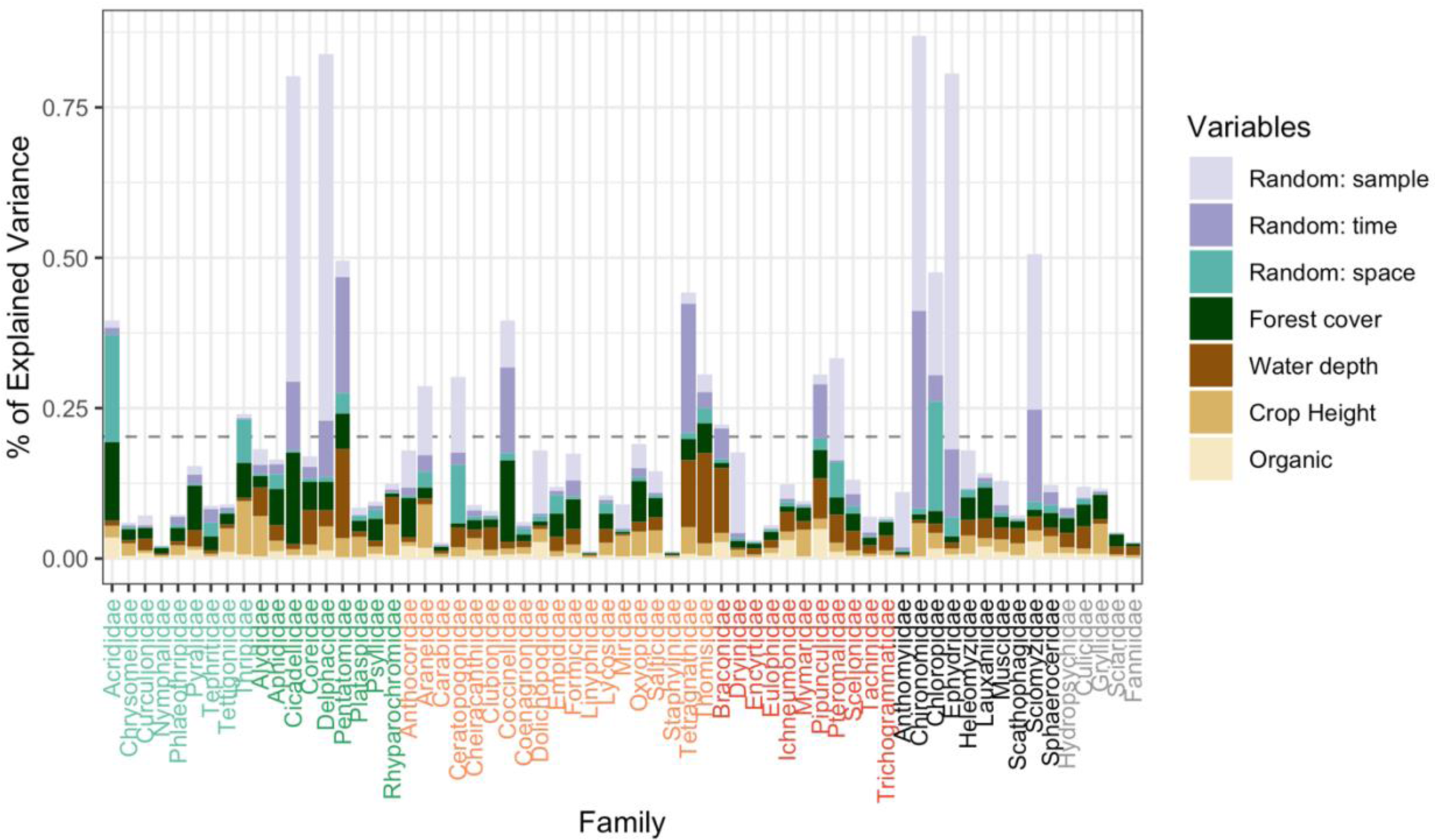
Variance partitioning of fixed and random effects of JSDM. Explained variance (%) of each species were adjusted by explanatory power, measured as Pseudo-R^2^. The dashed horizontal line indicates mean explained variance (0.20) across all species. Family names on the x-axis are arranged and color coded by trophic guild. (Light green = Leaf-chewer; Dark green = Sap-sucker; Orange = Predator; Red = Parasitoid; Black = Tourists; Grey = Filter-feeder, Scavenger, and Unknown).

### Family- and guild-specific responses to the environment

Results of species niches (β parameter estimates) show that there is large variation in species responses in terms of both sign and effect size (Fig. 4). Congruent with the results of variance partitioning, forest cover had the highest number of significant effects on species abundances, with the most effects being positive. Families that show the strongest response to forest cover were Gryllidae (β = 1.10; 90% CI: 0.219, 2.082) and Acrididae (β = 0.953; 90% CI: 0.293, 1.604). In contrast, effects of water depth were generally negative and had a lower number of significant effects. The family showing the strongest response to water depth were Fanniidae (β = -1.141; 90% CI: -2.076, -0.236) and Pentatomidae (β = - 0.942; 90% CI: -1.261, -0.638). Families showed less significant responses to crop height and organic farming but showed more variability in the sign of effects (Fig. 4). Families showing the strongest response to crop height were Gryllidae (β = -1.290; 90% CI: -1.898, - 0.727) and Miridae (β = -0.967; 90% CI: -1.36, -0.590). For organic farming, Acrididae (β = 0.956; 90% CI: 0.261, 1.647) and Pipunculidae (β = -0.844; 90% CI: -1.377, -0.337).

**Figure 4.**
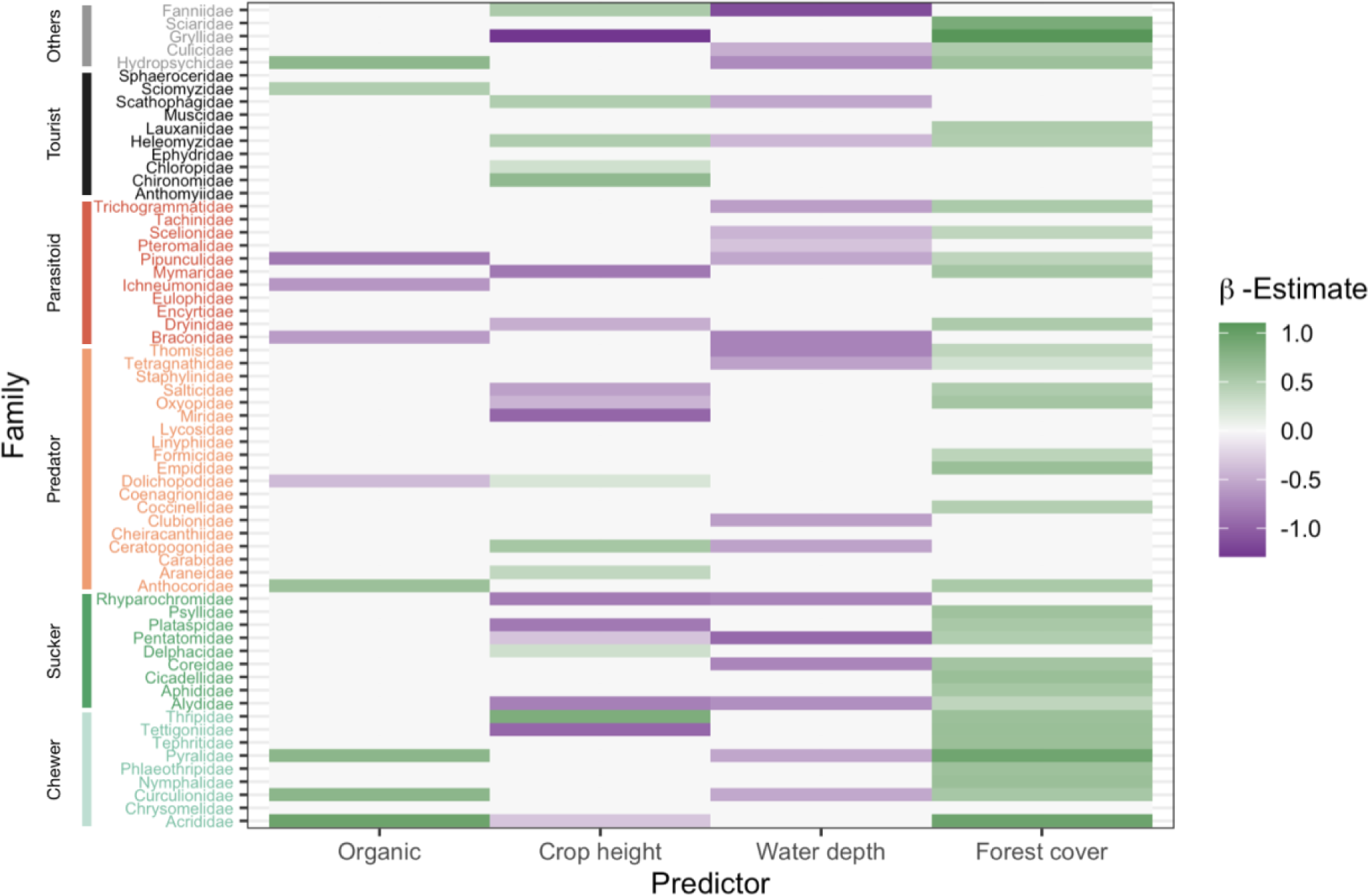
Heatmap summarizing estimated species responses (β) to environmental covariates. Non-significant parameter estimates (90% credible intervals that contain 0) are set to 0, significant estimates are set to its posterior mean. Family names are arranged and color coded by guild as described in Figure 3.

At the trophic guild level, guild explained 15.1% of the variation in species responses to environmental variables (Fig. S1). However, many of these parameters were not significantly different from 0 at the 90% significance level. In contrast, PERMANOVA results indicate that trophic guilds explained 33% of the variation in multidimensional niche space (mean β estimates regardless of significance; Fig. 5, Table S2). When projecting species responses in ordination space, the main axis of variation (PC1) was poorly correlated with the 4 environmental covariates and was qualitatively unable to distinguish between guilds (Fig. 5). In contrast PC2, although explaining less variation (26%), captured the distinction in niches between leaf chewers and parasitoids. Organic farming and water depth were the variables that show the most heterogeneity in species responses (indicated by the vector lengths; Fig. 5).

**Figure 5.**
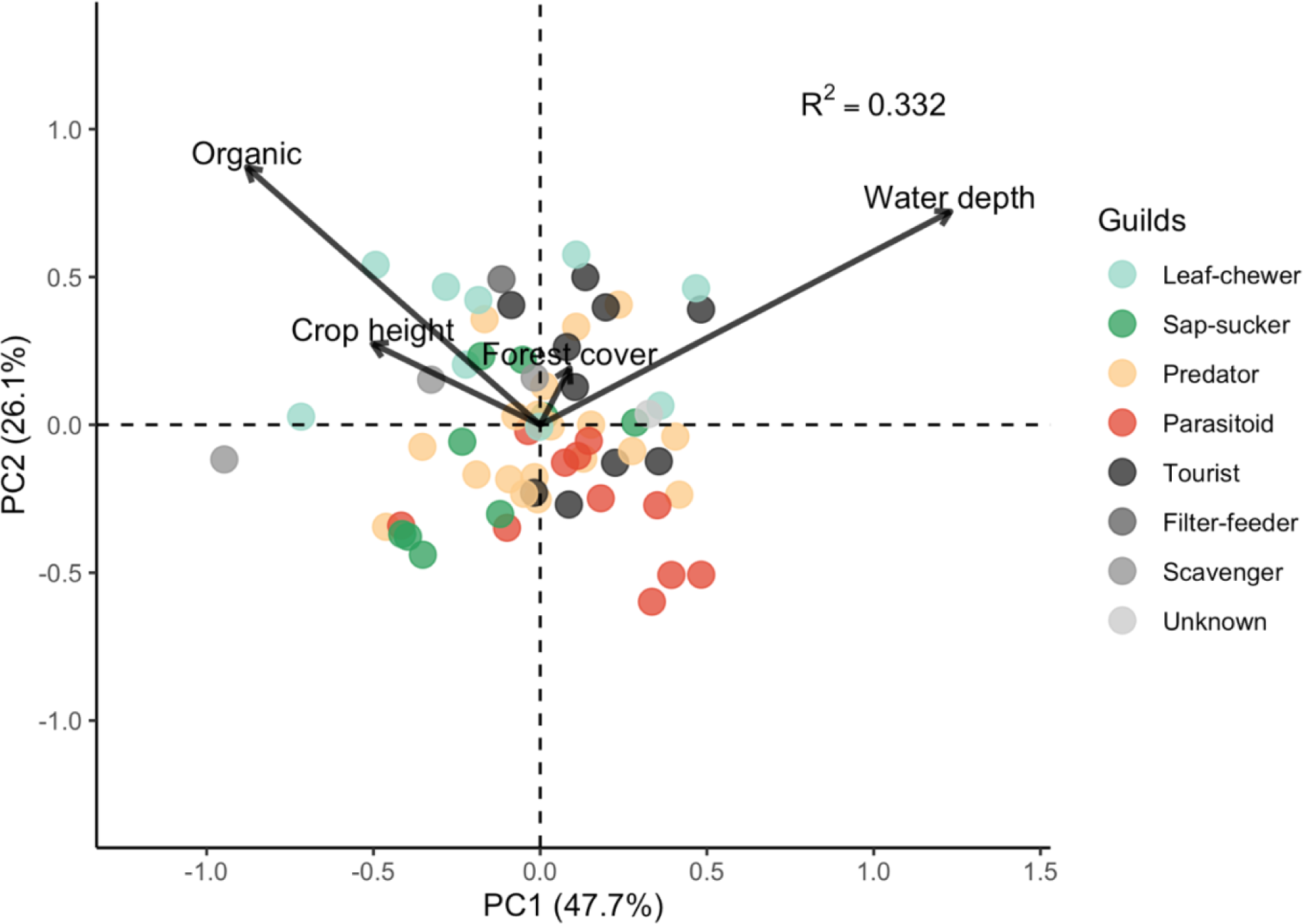
Principal component analysis (PCA) of mean family responses (β) by trophic guilds. Points indicate the posterior means of the 4 environmental covariates of each family, reduced to 2 major axes (PC1 & PC2) of variation. Vector arrows show the optimal responses of environmental covariates and their respective angular orientation represent correlations between them. Length of vector arrows represent the contribution of the covariate to the multidimensional variation of mean family responses. R^2^ value indicates explained variation of trophic guilds from PERMANOVA. Points are color coded by trophic guilds.

### Mean and variability of environmental effects across species

Analyses of the mean and variability of niches (β) across families show that the effect of environmental covariates vary in magnitude and sign but also vary in the degree of coherence among species (Fig. 6). Specifically, the “average species” responded positively to organic farming (mean = 0.162; 90% HDI: -0.0131, 0.322) and forest cover (mean = 0.397; 90% HDI: 0.280, 0.509) while responding negatively to crop height (mean = -0.068; 90% HDI: -0.162, 0.015) and water depth (mean = -0.311; 90% HDI: -0.420, 0.218). In terms of variability, responses were more variable to organic farming (mean = 0.532; 90% HDI: 0.415, 0.637) and crop height (mean = 0.491; 90% HDI: 0.414, 0.568) and less so for water depth (mean = 0.388; 90% HDI: 0.313, 0.461) and forest cover (mean = 0.359; 90% HDI: 0.268, 0.453).

**Figure 6.**
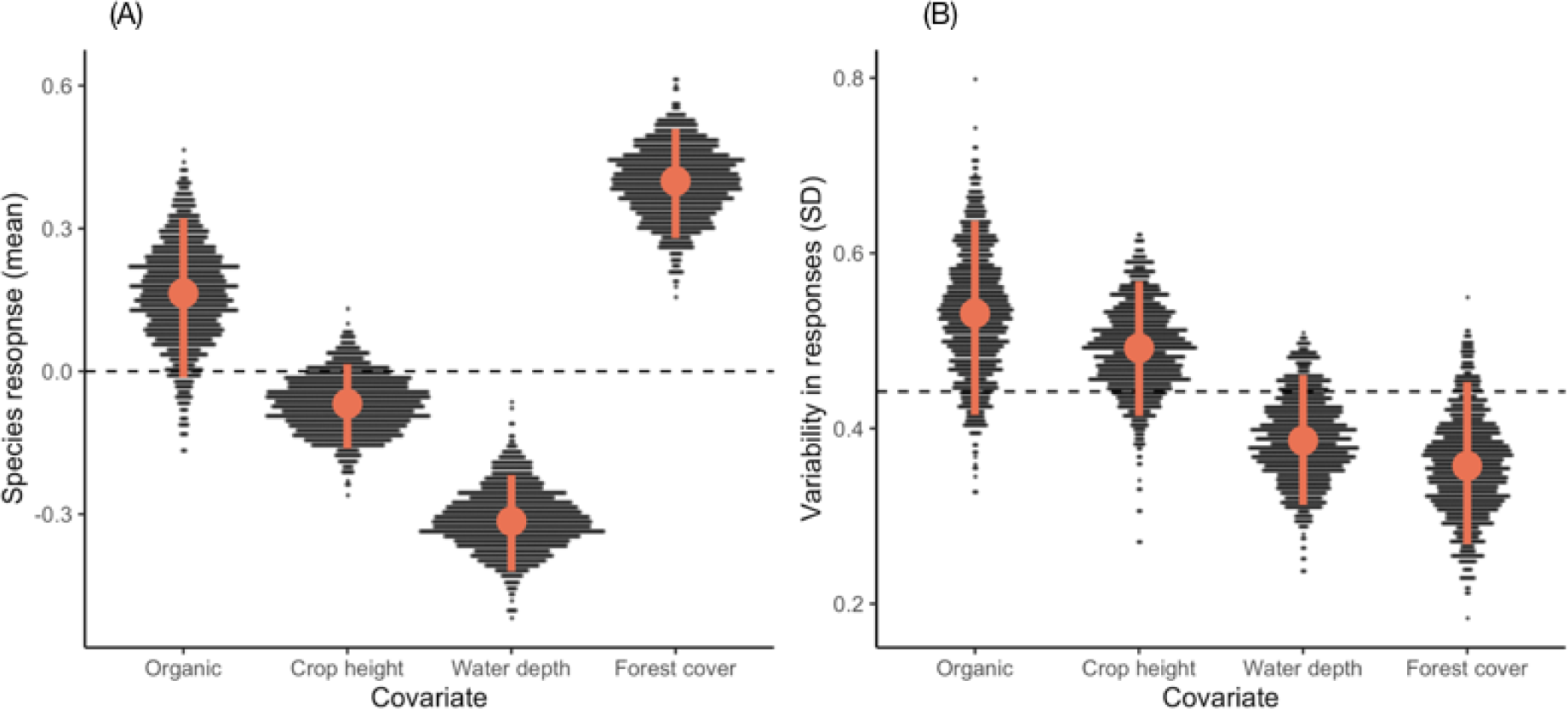
Sina plot showing the mean (A) and variability (B) of responses to environmental covariates across family. Data points plotted are derived from drawing 1000 random subsample from the 2 MCMC chains pooled together. Red point indicates mean of 1000 random subsamples while the error bar indicates the 90% highest density interval (HDI).

### Effects of space and time on species abundance and distribution

The effect of time but not space was prevalent in explaining arthropod species abundances of our system. The effect of space, or the extent of dispersal limitation in structuring species abundances, was minuscule (Fig. S2). Inspection of the two leading spatial latent variables, we see that their posterior distribution overlap 0, indicating that there is no residual variation of species abundances that can be explained by space that is independent of environmental covariates (Fig. S2A, B). This can be further verified by inspecting the residual variation captured by the η parameters plotted on the geographic coordinates of our samples (Fig. S2C, D). This indicates that dispersal limitation in our system is low, species are well-mixed across the landscape, and that variation in the distribution and abundance of species is driven by environmental conditions and stochastic processes. On the other hand, species abundances followed an obvious temporal trajectory which indicates a strong temporal turnover of species composition (Fig. S3). Specifically, the temporal scale of species turnover was estimated to be approximately 36 days (Fig. S3), which also corresponds approximately to the time it takes for the rice crop to enter its respective growth stages (i.e. tillering, flowering, and ripening).

### Effects of environment on community-level richness

At the community-level, we found that farm type had only a marginal effect on richness (χ^2^ = 3.224, df = 1, P = 0.064; Fig. 7) (Table 1). Community richness both decreased with crop height (χ^2^ = 15.899, df = 1, P < 0.001) and water depth (χ^2^ = 23.290, df = 1, P < 0.001). In Contrast, community richness increased with surrounding forest cover (χ^2^ = 83.136, df = 1, P < 0.001). With respect to model fit and importance of fixed effects, measured as the explained family richness, the most important factor in explaining richness was forest cover (R^2^= 0.278), followed by water depth (R^2^= 0.121), crop height (R^2^= 0.058), and farm type (R^2^= 0.012).

**Figure 7.**
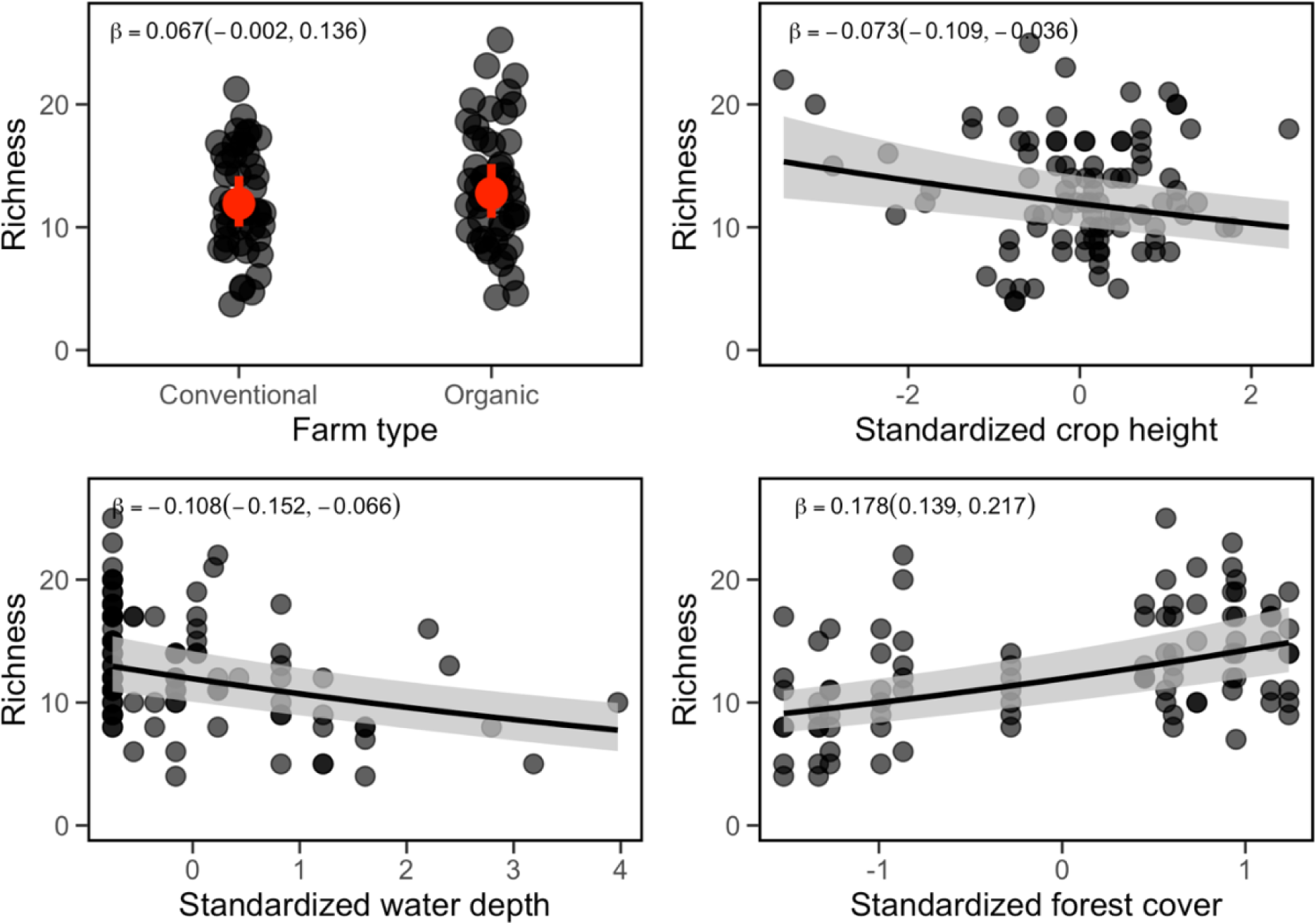
Partial effects of environmental covariates on predicted community richness. Points indicate the community richness of empirical samples (N = 94) predicted by JSDM taken as the posterior median. Farm type (A) showed no significant effect on predicted community richness. Community richness was negatively correlated with crop height (B) and water depth (C). Forest cover (C) showed a positive correlation with community richness. Red points (A) and fitted lines (B-D) indicate the predicted values from GLMM respectively. Error bars (A) and confidence bands (B-D) show 95% confidence interval of predictions.

**Table 1.** Results of generalized linear-mixed effect model of sample species richness predicted by JSDM weighted by posterior SD. Significance of predictors were assessed by likelihood ratio tests.

| **Predictor** | **$\beta$** | **SE** | **$\chi^2$** | **$P$** |
| --- | --- | --- | --- | --- |
| Intercept | 2.480 | 0.086 |  |  |
| Organic | 0.067 | 0.035 | 3.225 | 0.064 |
| Crop height | -0.073 | 0.019 | 15.899 | <0.001 |
| Water depth | -0.108 | 0.022 | 23.290 | <0.001 |
| Forest cover | 0.178 | 0.020 | 83.136 | <0.001 |

## Discussion

### 1. How do species differ in their responses to local and landscape factors?

The effects of local variables on farmland arthropod biodiversity were highly variable across species and covariates. Despite common belief, the effects of organic farming was largely negligible at the community-level, albeit showing significant positive and negative effects on a few selected families. Interestingly, of the few families that show significant responses to organic farming, leaf-chewer herbivores tend to show positive responses while parasitoids showed negative responses (Fig. 2). Although this finding may be discouraging for farmers, herbivores in the leaf-chewer guild are of less importance to the rice crop yield compared to sap-sucker herbivores, which contain damaging pests such as planthopper, leafhoppers, and rice seed bugs (Bottrell & Schoenly, 2012). More importantly, our finding is converse to the conclusions of a previous meta-analysis (Bengtsson et al., 2005), suggesting that generalities on the effects of organic farming on species diversity need to be made with caution. Whether rice agroecosystems are an exception to these general claims remains an unanswered and important question to address due to their underrepresentation in meta-analyses (Settele & Settle, 2018).

Among environmental covariates, the effect of crop height was the most enigmatic. For example, crop height may provide additional structural complexity and available niches that should maintain more speciose communities. However, the effect of crop height was variable across species, with almost equal proportions of positive, negative and non-significant effects. In addition to the structural effects of height itself, it is possible that correlated traits such as plant quality and defense may also influence the growth and abundance of herbivores. Interestingly, two dominant pest species of rice in the Delphacidae and Thripidae families, showed positive responses to crop height (Fig. 4). Being herbivores highly specialized to feed on rice crops, it is likely that the effect of crop height is related to plant nutritional quality because specialized species have been shown to be less sensitive to variable plant quality (Lin et al., 2020). However, more work is required to confirm the presence of height-quality correlations and determine whether it is the mean or variability in plant quality that affects herbivore performance (Wetzel et al., 2016).

Our study provides, to our knowledge, the first documentation of the effect of water depth on paddy field arthropods. Surprisingly, water depth had an overall negative effect and was relatively consistent across species. Given that the primary use of water in rice agriculture is to limit weed growth, our results suggest a tradeoff between weed control and diversity maintenance. Additionally, as agricultural wetlands provide habitat for many waterfowls (Czech & Parsons, 2002), it is also likely that arthropod abundances are reduced due to either direct consumption or disturbance by bird activity.

One of the defining features of agroecosystems is that factors of agricultural intensification can occur on local and landscape scales (Tscharntke et al., 2005). In our study, forest cover served only as an index — rather than a direct measure — of landscape complexity, yet, it was the strongest and least variable predictor of species abundances. Numerous mechanisms may explain this finding. For instance, because forest cover is a percentage variable, it is possible that negatively correlated variables such as human development, but not forest cover *per se*, may be the actual factor that affects species abundances. However, the effect of forest cover on biodiversity is a well-documented phenomenon in agroecosystems as it provides necessary habitats, areas for refuge (Corbett & Rosenheim, 1996; Kormann et al., 2015), additional resources (Sutter et al., 2017), and structural complexity (MacArthur, 1958) needed for maintaining species diversity. Moreover, higher primary production of the landscape may also contribute to higher consumer diversity (Huston & Wolverton, 2009).

The relative importance of local and landscape factors on biodiversity ultimately depends on the species of interest, their life-history traits, and habitat requirements. For example, arthropods in particular, contain many species that require non-crop vegetation and aquatic habitats to complete stages of their life cycle (Bugg et al., 2008). A diverse landscape that contains various ecosystem types will therefore be critical for maintaining their population persistence. However, the distinct features of the landscape are likely to contribute differentially to species persistence. Moreover, dispersal is also not the only type of organismal movement. A promising road forward is to integrate the meta-ecosystem framework (Gounand et al., 2018; Loreau et al., 2003) and incorporate more types of organismal movement (e.g. foraging, life-cycle, migration), to understand how spatial flows of material and energy across ecosystem boundaries may contribute to the structure and persistence of ecological communities.

### 2. Can trophic guilds be used as a good proxy of species responses to environmental variables?

In many cases, species differ in their relevance to conservation and/or agriculture. As such, there is a practical need for identifying the relative importance and the effects of environmental variables on particular species. Taking a species-centric approach using JSDM, we show that species in our system do indeed show large variation in how they respond to (i.e. β) and the relative importance of (i.e. R^2^) environmental covariates. Moreover, heterogeneity of species responses also differed substantially among covariates. These findings suggest that land-use practices that aim to increase biological diversity to enhance ecosystem services must do so with caution because richness increases may arise from a different set of species. Importantly, due to the low coherence of species responses within trophic guilds (Fig. 3, 4) our study refutes the common assumption that broad functional groups serve as good proxies for species responses and highlight the limitations of studies that infer community dynamics and trophic interactions using these broad classifications without experimental manipulations (Barrion et al., 1994; Dominik et al., 2018; Drechsler & Settele, 2001; Heong et al., 1992; Macfadyen et al., 2015).

Interestingly, the large variation in the importance of environmental covariates suggests the presence of core-transient dynamics (Hanski, 1982; Holt & Gaines, 1992; Pulliam, 1988) which is expected to be prevalent in agroecosystems due to their ephemeral nature (Vandermeer et al., 2010). Specifically, core species respond more strongly to environmental covariates while the reverse is true of transient species. Presence of these dynamics support the notion that communities do not respond as a unit and that the relative strength of deterministic and stochastic processes may not be generalizable across all species (Coyle et al., 2013; Umaña et al., 2017; White & Hurlbert, 2010). These dynamics are ecologically interesting in their own right but also have important conservation implications as they may serve as a mechanism for regional species coexistence (Leibold et al., 2004; Levins & Culver, 1971) but also dictate the extent to which diversity can enhance ecosystem functions or services (Leibold et al., 2017). Thus, we suggest that future work can be designed to investigate which components across the landscape act as source pools and to delineate the service providers among source-sink species to ensure species persistence and the provisioning of ecosystem services.

### 3. How do species-specific responses scale-up to structure arthropod biodiversity (i.e. species richness)?

By incorporating species-level responses, our study reveals the different pathways in which community richness can arise. For instance, although water depth and crop height were both negatively correlated with community richness, their effects on individual species were very different. Specifically, the negative effect of water depth was relatively consistent across species while being highly variable for crop height (Fig. 4 & 7). Similarly, although organic farming had no effect on family richness, notable effects on the abundances of a few families were still found. Without information on the responses of the constituents of biodiversity (i.e. individual taxa), false conclusion on the effects of environmental variables may be drawn.

## Conclusion

By taking a species-centric approach, our study highlights the limitations and opportunities of local and landscape management for biodiversity conservation. For instance, local land-use practices such as organic farming, crop attributes and weed management may show weaker impacts on arthropod diversity overall in comparisons to landscape complexity, but their effects may still be substantial for specific species. Furthermore, our results indicate high heterogeneity in species responses, even within broad functional groups (i.e., trophic guilds). This finding warrants the need for taking species-centric approaches (Betts et al., 2014; Frishkoff & Karp, 2019) and to take cautionary measures when quantifying biodiversity change based on broad functional group classifications. Our analysis of the residual variation in species abundances indicate a strong turnover in species composition over time — as is expected with paddy field arthropods (Settle et al 1996; Schoenly et al 1996) — but not in space, which suggests that the lack of dispersal limitation in structuring communities. Lastly, by scaling up biodiversity from individual species responses, we reveal the relative importance of environmental variables of agriculture and the pathways in which they can structure paddy field arthropod communities.

## Declaration of Competing Interest

The authors declare no competing interests.

## Data availability

Data will be made available upon request.

## Acknowledgements

We thank the Miaoli District Agricultural Research and Extension Station, in particular, Hung-Ju Chen for logistical support. We also thank Nana Zou for assisting with creating land-use maps as well as the private landowners for access to their farms. This research was funded by Council of Agriculture, Executive Yuan, Taiwan (xxxx).

## Author contributions

J.-A. Ou and C.-K. Ho conceived of the study. J.-A. Ou collected geospatial data, conducted statistical tests, and wrote the manuscript with input from C.-K. Ho. C.-L. Huang collected and identified arthropod data. C.-W. Tsai and C.-K. Ho secured funding.

## Notes

### Competing Interest Statement

The authors have declared no competing interest.

### Summary of Updates

Figure legend; Author list updated

