## Supplementary material for "Effects of local and landscape factors on paddy field arthropod diversity: from species to communities"

Table S1. Guild assignments of the families analyzed.

| Guild | Family |
| --- | --- |
| Filter-feeder | Hydropsychidae |
| Leaf-chewer | Acrididae, Chrysomelidae, Curculionidae, Nymphalidae, Phlaeothripidae, Pyralidae, Tephritidae, Tettigoniidae, Thripidae |
| Parasitoid | Braconidae, Dryinidae, Encyrtidae, Eulophidae, Ichneumonidae, Mymaridae, Pipunculidae, Pteromalidae, Scelionidae, Tachinidae, Trichogrammatidae |
| Predator | Anthocoridae, Araneidae, Carabidae, Ceratopogonidae, Cheiracanthiidae, Clubionidae, Coccinellidae, Coenagrionidae, Dolichopodidae, Empididae, Formicidae, Linyphiidae, Lycosidae, Miridae, Oxyopidae, Salticidae, Staphylinidae, Tetragnathidae, Thomisidae |
| Sap-sucker | Alydidae, Aphididae, Cicadellidae, Coreidae, Delphacidae, Pentatomidae, Plataspidae, Psyllidae, Rhyparochromidae |
| Scavenger | Culicidae, Gryllidae, Sciaridae |
| Tourist | Anthomyiidae, Chironomidae, Chloropidae, Ephydridae, Heleomyzidae, Lauxaniidae, Muscidae, Scathophagidae, Sciomyzidae, Sphaeroceridae |
| Unknown | Fanniidae |

Table S2. Results of PERMANOVA testing the effects of species guilds on the posterior mean responses to multiple environmental covariates.

|  | **Df** | **Sum of Squares** | **Mean Sum of Squares** | ***F*** | **R^2^** | ***P*** |
| --- | --- | --- | --- | --- | --- | --- |
| Guilds | 7 | 9.440 | 1.349 | 3.902 | 0.332 | 0.001 |
| Residuals | 55 | 19.008 | 0.346 |  | 0.668 |  |
| Total | 62 | 28.448 |  |  | 1.000 |  |

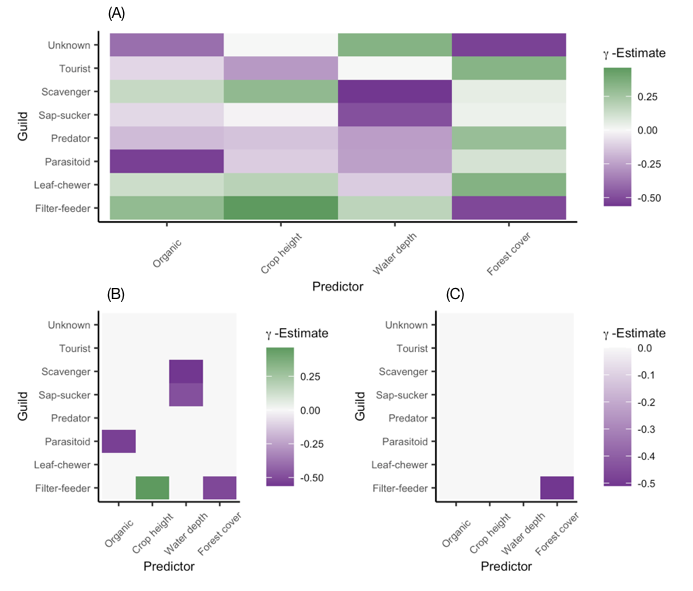

Figure S1. Heatmap summarizing estimates responses of trophic guild (γ) to environmental covariates. Panel (A) shows the mean responses regardless of significance, panel (B) shows significance at 80%, and panel (C) show significance at the 90% level. Overall, trophic guild explained 15.1% of the variation in family-specific responses to covariates.

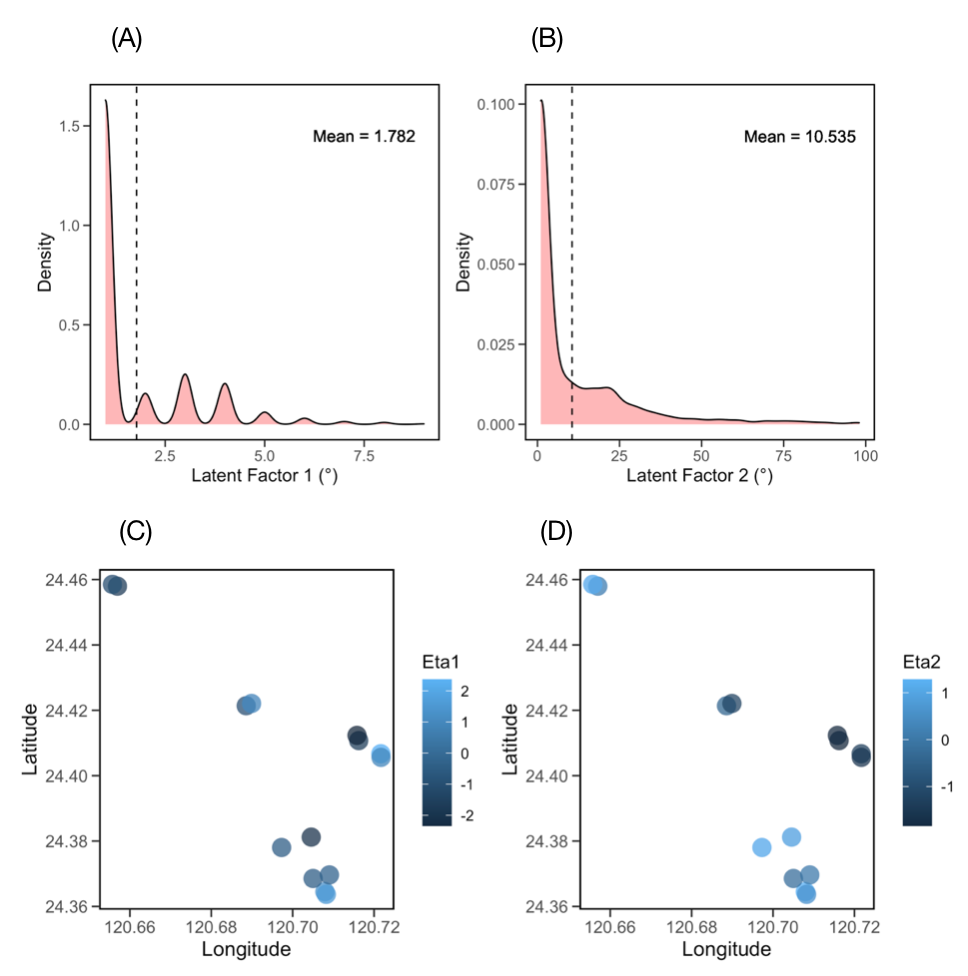

Figure S2. Estimated spatial scale and spatial residual variation of abundances plotted in space. Spatial latent factors estimate the Euclidean distance between geographic coordinates that explains variation in abundances among samples. Both leading latent factors were overlap 0, indicating the lack of dispersal limitation. Eta (η) estimates the residual abundance variation for each site. No qualitative pattern can be observed by plotting Eta in space, consistent with the lack of spatial autocorrelation.

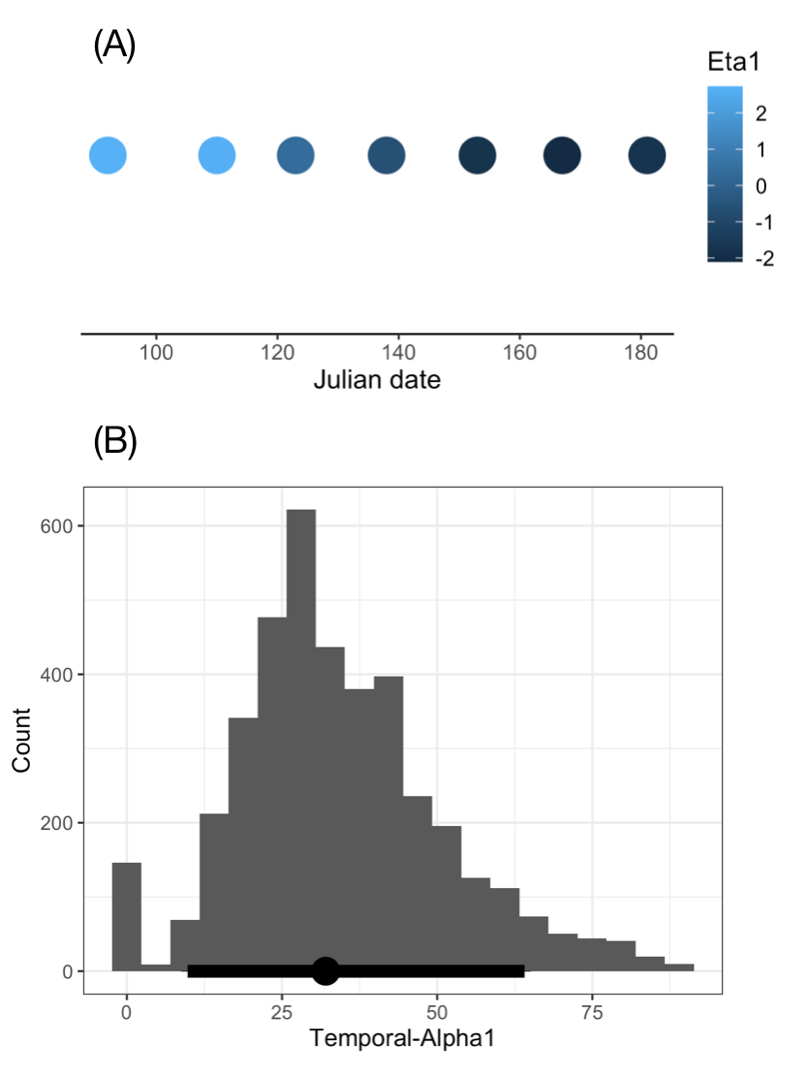

Figure S3. Temporal residual variation of abundances and estimated temporal scale. Residual variance of species abundances changes over time (A), indicating turnover in species composition. Estimated temporal scale is 36 days (B).
